# New Pathways of Mutational Change in SARS-CoV-2 Proteomes Involve Regions of Intrinsic Disorder Important for Virus Replication and Release

**DOI:** 10.1101/2020.07.31.231472

**Authors:** Tre Tomaszewski, Ryan S. DeVries, Mengyi Dong, Gitanshu Bhatia, Miles D. Norsworthy, Xuying Zheng, Gustavo Caetano-Anolles

## Abstract

The massive worldwide spread of the SARS-CoV-2 virus is fueling the COVID-19 pandemic. Since the first whole-genome sequence was published in January 2020, a growing database of tens of thousands of viral genomes has been constructed. This offers opportunities to study pathways of molecular change in the expanding viral population that can help identify molecular culprits of virulence and virus spread. Here we investigate the genomic accumulation of mutations at various time points of the early pandemic to identify changes in mutationally highly active genomic regions that are occurring worldwide. We used the Wuhan NC_045512.2 sequence as a reference and sampled 15,342 indexed sequences from GISAID, translating them into proteins and grouping them by month of deposition. The per-position amino acid frequencies and Shannon entropies of the coding sequences were calculated for each month, and a map of intrinsic disorder regions and binding sites was generated. The analysis revealed dominant variants, most of which were located in loop regions and on the surface of the proteins. Mutation entropy decreased between March and April of 2020 after steady increases at several sites, including the D614G mutation site of the spike (S) protein that was previously found associated with higher case fatality rates and at sites of the NSP 12 polymerase and the NSP13 helicase proteins. Notable expanding mutations include R203K and G204R of the nucleocapsid (N) protein inter-domain linker region and G251V of the viroporin encoded by ORF3a between March and April. The regions spanning these mutations exhibited significant intrinsic disorder, which was enhanced and decreased by the N-protein and viroporin 3a protein mutations, respectively. These results predict an ongoing mutational shift from the spike and replication complex to other regions, especially to encoded molecules known to represent major β-interferon antagonists. The study provides valuable information for therapeutics and vaccine design, as well as insight into mutation tendencies that could facilitate preventive control.

## Introduction

The first case of ‘coronavirus disease 2019’ (COVID-19) was identified in the Chinese city of Wuhan in December 2019. Since then, the novel virus has rapidly spread to 188 countries and territories, infecting more than 15 million people and causing over 0.6 million deaths^1,2^. COVID-19 patients develop a ‘severe acute respiratory syndrome’ analogous to that of the 2002-2003 SARS epidemic that spread to 23 countries, infected ∼8,000, and killed 774 people. The COVID-19 virus was named SARS-CoV-2 by the WHO, and is the seventh coronavirus known to infect humans^2,3^. Currently, there are no vaccines or antiviral drugs capable of preventing or treating human infections^4^.

SARS-CoV-2 belongs to the *Betacoronavirus* genus of the *Coronaviridae* family, a group of related enveloped positive-sense single stranded RNA viruses that infect both mammals and birds. In chickens, viruses cause upper respiratory tract diseases, while in cows and pigs they cause diarrhea. In humans, viruses can cause respiratory tract infections that can range from mild to lethal. For example, the well-known SARS-CoV-2-related SARS-CoV-1 and MERS-CoV coronaviruses caused severe diseases in the zoonotic outbreaks of 2002-2003 and 2012-onwards, respectively^5–8^. Compared with its siblings, SARS-CoV-2 spreads more rapidly, infects more people, and shows a much lower fatality rate^1,9,10^.

SARS-CoV-2 has a ∼30 kb genome, which was first reported on January 5, 2020^8^. The genome encodes both structural and non-structural proteins. The leader sequence and ORF1ab encode non-structural proteins (NSPs) functioning in replication and transcription^11^. Together with accessory proteins, the structural proteins are encoded in the downstream regions of the genome. They include the spike (S) protein of the viral ‘corona’, the envelope (E) protein, the membrane (M) protein, and the RNA-binding nucleocapsid (N) protein. Coronaviruses infect human cells by using the homotrimeric spike glycoprotein, known as S-protein, to bind the angiotensin-converting enzyme 2 (ACE2) receptor, which is located in the epithelia of the lung and small intestine of humans^12,13^. SARS-CoV-2 has an optimized receptor-binding domain (RBD) that binds with high affinity to ACE2 in human and animals^14^. High receptor homology was supported by a series of structural and biochemical studies^14–19^. When compared with SARS-CoV, S-protein amino acids at positions 455 (leucine) and 486 (phenylalanine) on SARS-CoV-2 showed an enhanced interaction with hot spot 31 and viral binding to human ACE2^16^. These studies tested the origin of the virus, challenging the suggestion that SARS-CoV-2 represents an artificially designed manipulation. Instead, it likely arose as a novel recombinant virus transmitted from both horseshoe bats and pangolins^20^. Thus, transmission to and among humans appears a result of natural selection^14–16,21^.

Viral genome sequence data has been collected in real-time from COVID-19 patients at significant pace. By May 7, 2020, the GISAID database (https://www.gisaid.org/) gathered 15,366 full sequences of human coronavirus. This data provided opportunities to explore when, where and how mutations happen within the SARS-CoV-2 genome. The majority of mutational studies thus far have mainly focused on the S-protein. These studies revealed that the RBD is the most variable region, with several RBD amino acids showing critical ACE2 receptor binding functions^8,14–16,22^. S-protein mutations adjusted the binding efficiently to its human receptor. In contrast, mutations in other genomic regions have been rarely studied. Besides the notable S-protein, mutations on the N-protein that makes the nucleocapsid could alter virulence and virus spread. The N-protein (50 kDa) is the most abundant in both viruses and virus-infected cells, playing multiple roles in the replication and transcription of the virus, as well as in the assembly of the viral genome^23^. The N-protein binds to the viral RNA genome at its N-terminal end, forming a ribonucleoprotein complex that plays essential roles in maintaining a functional RNA conformation^24^. Due to it being highly immunogenic and possessing a normally conserved amino acid sequence, the SARS-Cov-2 N-protein is an optimal target for both diagnostic assays and vaccine formulations^23,25^.

Here we study pathways of molecular change in the genomes of the expanding SARS-CoV-2 population that can help identify molecular culprits of virulence and virus spread. Using the mutational entropy of nucleotide and amino acid sequences as a measure of molecular diversity over time, we study the evolutionary trajectory of crucial viral proteins such as the S- and N-proteins and several NSP and accessory proteins as we trace mutational changes in the sequences of 12,606 SARS-CoV-2 genomes. Remarkably, we find that while virulence-associated mutations in the S-protein are becoming fixed in time, nucleotide changes in the N-protein and the viroporin protein 3a that we find are associated with protein intrinsic disorder are establishing themselves as more prominent. Here we explore their significance.

## Materials and Methods

### Data

The SARS-CoV-2 reference sequence, accession NC_045512.2 (version March 30, 2020; previously ‘Wuhan seafood market pneumonia virus’) was obtained from the NCBI Virus repository on May 4, 2020^8^. A total of 15,366 sequences were acquired on May 7, 2020 from GISAID^26^ and its initiative’s platform EpiCoV (see Supplementary Table 1 for sequence information). Metadata was acquired from NextStrain^27^. The sequence data and metadata were mapped using the GISAID EPI_ISL number as the primary key. The metadata revealed 24 non-human host sequences, which were removed. Any primary keys not appearing in both the metadata and sequences were also removed.

**Table 1.**
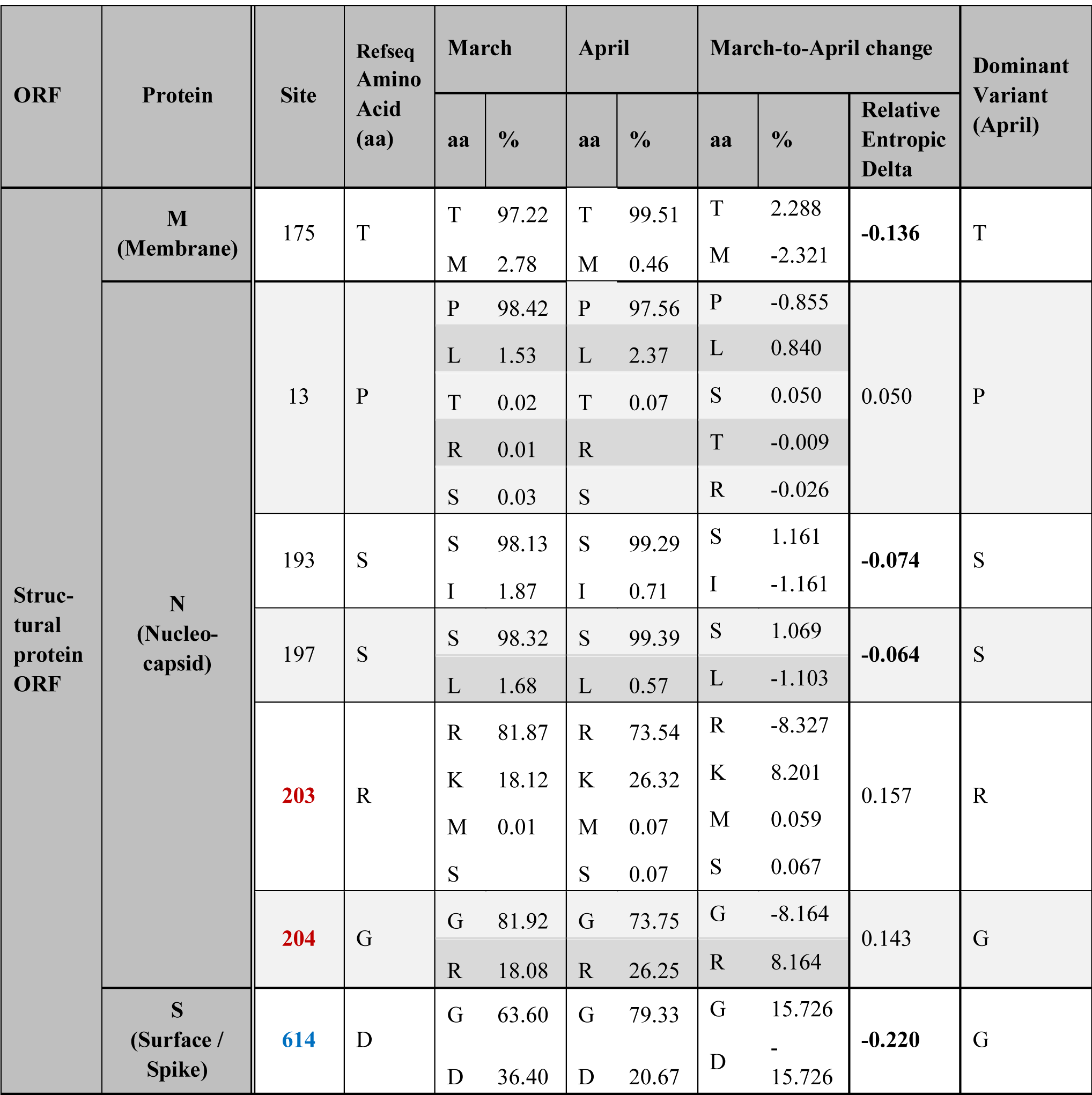
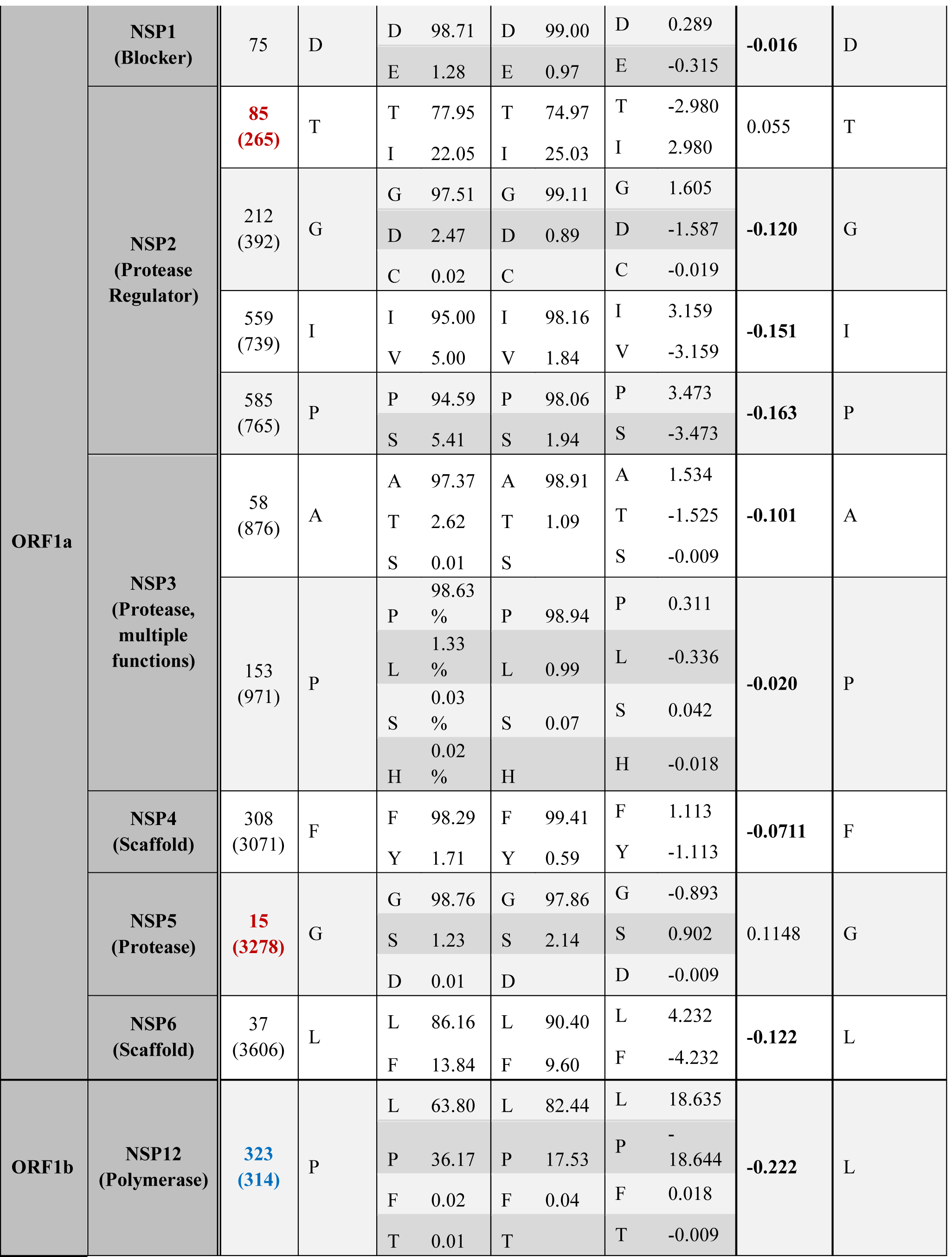
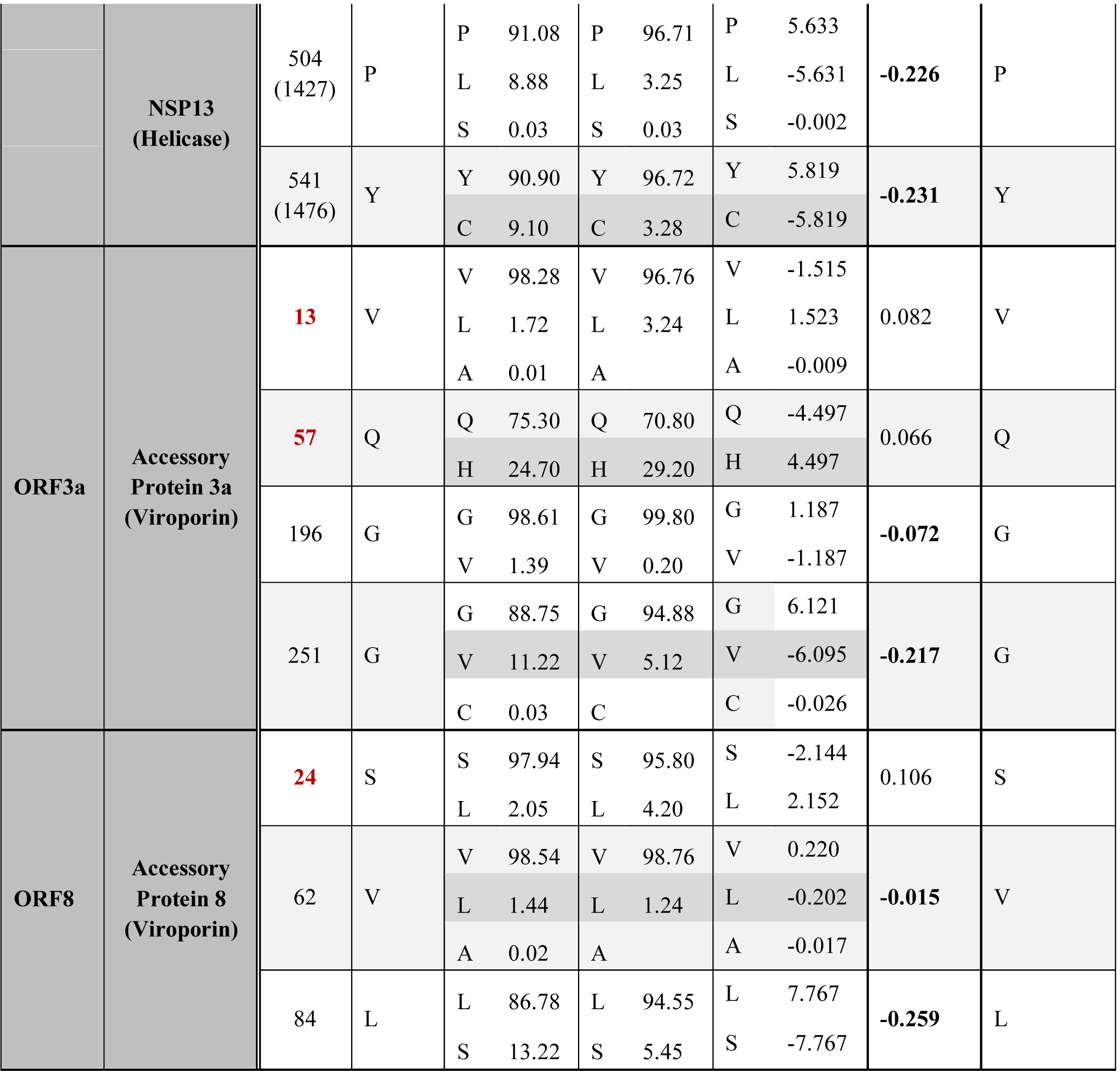
Amino acid sites with major entropy changes between the months of March and April. Site numbers labeled in blue indicate locations where an amino acid variant is currently dominant but differs from the RefSeq entry. Site numbers labeled in red indicate locations where a variant is currently non-dominant but is ascendant by >1% between March and April. Relative entropic delta values in bold indicate entropic decreases.

The remaining 15,342 sequences were then aligned with the reference sequence, removing any gaps caused by the initial inclusion non-human host sequences. The head (<g.266) and tail (>g.29,684) sections were removed, as these sites lie outside all known coding regions and are highly variable in composition and length across sequences, providing little salient information for protein analysis. The sequences were then split into coding region sequences (CDS) corresponding to NSPs, structural and accessory protein regions, as listed by the reference sequence GenBank document. These sub-sequences were translated into proteins using BioPython^28,29^, requiring replacement of all gap characters in the nucleotide sequence with ‘N’ characters, and for additional ‘N’ characters to be added to the end if the end contained a split codon. The protein sequences were also categorized by month of collection, using the NextStrain metadata. The complete data workflow is summarized in Figure 1. The blue sections are the main workflow steps, while the annotations in grey show details or notes about each section.

**Figure 1.**
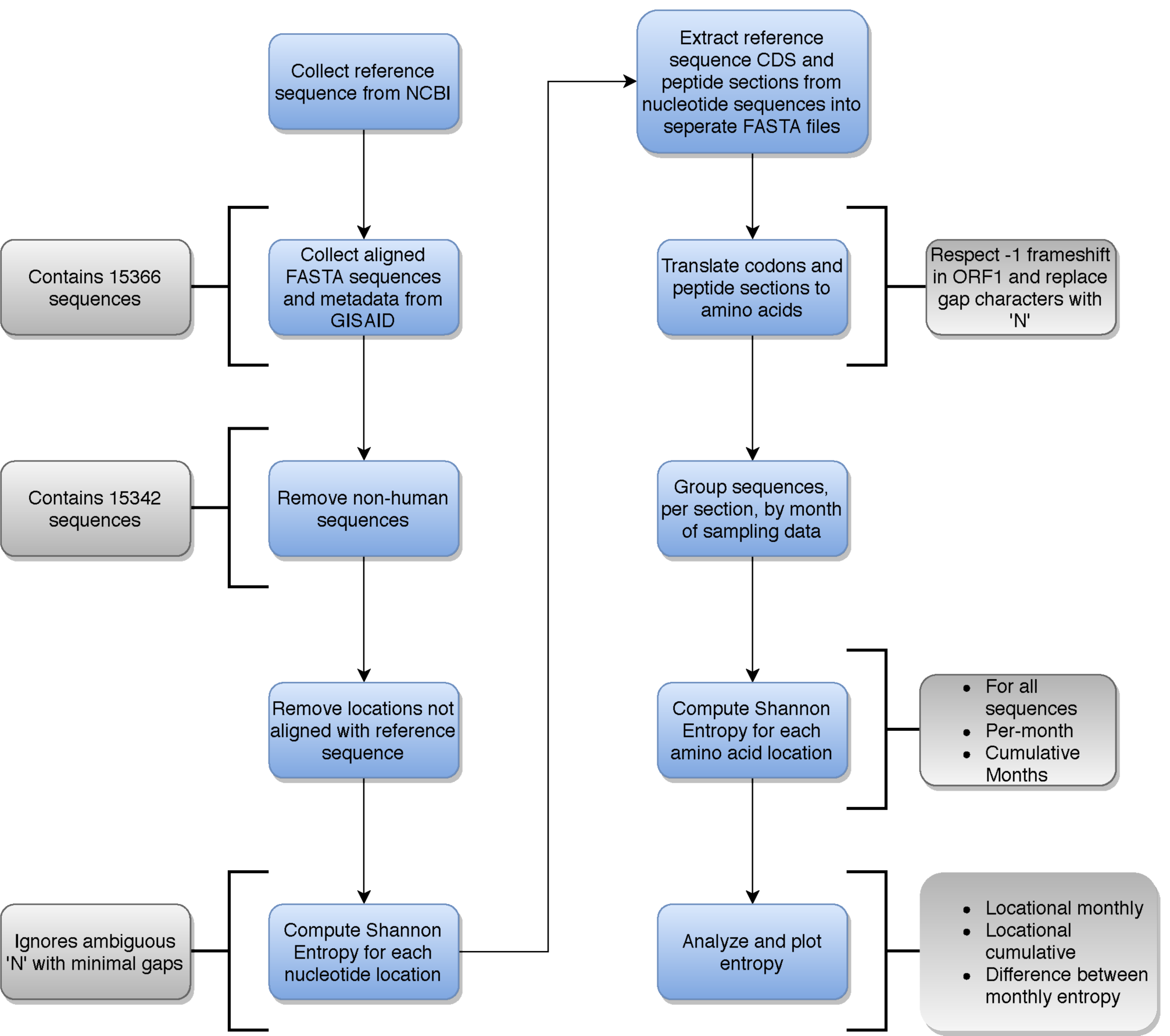
General data workflow of the analysis of SARS-CoV-2 genomes, including a breakdown of details that occur during each step. Main steps are indicated in blue, while details per step are indicated in gray.

### Analysis

The per-position amino acid frequencies and Shannon entropies were calculated across the CDS for each sampled month of the initial period of the pandemic (December through April). The number of genomes sampled (% total) increased in time, 18 genomes (0.1%) in December, 390 in January (2.5%), 612 in February (4.0%), 11,407 in March (74.4%) and 2,915 in April (19.0%). While the sample count per month varied widely, the difference should not affect the comparisons of per-location proportionality. To confirm the March dataset was not introducing noise due to the disproportionate size, the original entropy calculation as well as the entropies of 10 randomly sampled March subsets of 2,915 sequences (April’s sequence count) were compared using cosine similarity. The mean similarity between the original March entropy and the sampled entropy arrays was 0.997 (min: 0.9964, max: 0.9973). Likewise, 10 entropic samples of 612 March sequences (February’s count, 5.4% compared to March) had a mean similarity of 0.9851 (min: 0.9825, max: 0.9875).

Characters not included in the IUPAC standard protein alphabet (i.e., B, Z, J, U, O, and X) were ignored. This was justified through reasoning that (a) the amount of non-standard characters per sequence-set was negligible, and (b) the independent nature of the entropy of each location removes the chance of a cascading error. This exclusion also permitted the use of all obtained sequences, as sequences with high proportions of ambiguous characters would only contribute information if a valid amino acid were present at the analyzed location.

Calculation of pairwise distance metrics used a subset of distinct sequences per protein, allowing computation in reasonable time while maintaining a global view of the dataset, as all identical sequences will share pairwise distances. Sequences containing more than 5% ambiguous characters per protein were removed to further reduce the set and remove excess noise. Identical sequences were discovered through comparing hashes (SHA256) of the remaining sequence strings, producing a subset of distinct representative sequences. Pairwise distance calculations were performed using a BLOSUM80 (Block Substitution Matrix 80%) model^30^. This model was chosen based on the similarity between sequences was expected to be ≥80% as the set was a pre-made multiple sequence alignment containing *variants* of SARS-CoV-2 taken in a 5-month span. It should be noted that the BLOSUM models are not considered ideal for intrinsically disordered sequences^31^, but the pre-alignment, specific similarity of the sequences in the set, and the intended goal of analysis was not considered a severe detriment to the performance or outcome.

Downstream analyses included performing exploratory data analysis with principal component analysis (PCA), tracing mutations in crystallographic models, exploring if they fell in conserved or variable regions of the molecules, and analyzing of intrinsic disorder and binding potential in the proteins of the entire proteome. Dimensionality reduction was conducted using classical PCA on the distance matrices, capturing the landscape of observed distinct sequence variants. Mutations were traced onto published crystallographic models, or in their absence, i-Tasser *de novo* inferences^32^ downloaded from https://zhanglab.ccmb.med.umich.edu/COVID-19/. When possible, these inferences were later confirmed by structural alignments to regions that are structurally known. Structural neighborhoods were analyzed using DALI and sequence and structural conservation were mapped onto 3-dimensional models^33^. Finally, explored intrinsic disorder in the proteins of the SARS-CoV-2 proteomes. We categorized sequences as ‘highly structured’ (0-10% of the total protein length is unstructured), ‘moderately unstructured’ (10-30% of the protein is unstructured) and ‘highly unstructured’ (30-100% of the protein is unstructured) following the classification of Gsponer et al.^34^. A map of intrinsic disorder regions and binding sites within each of the reference sequence coding regions were produced using IUPred2A^35,36^. In addition, the Anchor2 algorithm implemented in IUPred was used to predict binding regions of viral protein based on the amino acid sequence. The algorithm is designed to identify segments in disordered regions that have the capability to gain energy by interacting with a globular partner protein. Anchor2 is based on the following three properties: (i) ensuring a residue belongs to a long disordered region and filters out globular domains, (ii) ensuring that the residue is not able to form enough favorable contacts with its own local sequential neighbors to fold, and (iii) ensuring there would be an energy gain upon interaction by testing the feasibility that a given residue can form favorable interactions with globular proteins upon binding^27,37^.

## Results

### Entropic Variation

Informational entropy describes the amount of variation in discrete per-location nucleotide or amino acid composition data. Here we study the evolutionary diversification of SARS-CoV-2 with an entropy-based strategy that quantifies diversity and selection in viral populations with two independent state variables^38^. First, we focus on mutational entropy as a measure of molecular biodiversity of the SARS-CoV-2 proteome. Mutational entropy is maximal when all amino acids in an amino acid site of the viral population are equally represented. Entropy is zero when only one amino acid overtakes that site. Increases in entropy imply dilution of mutations in the viral population while decreases signal fixation of those mutations. Second, we then consider a relative measure of entropy, ‘relative entropy delta’, which measures selection pressure by comparing viral diversity at two different time points of the pandemic and identifying entropy reversal trends suggestive of fitness advantage unfolding in time-space or entropy expansions suggesting other evolutionary forces are at play.

An initial analysis of mutational entropy at nucleotide level (Figure 2A) led to an evaluation of entropy at amino acid level (Figure 2B). Analyses identified several genomic sites of high protein entropy during at least one of the initial months of the pandemic. These were located for example in ORF1a, ORF1b, N, M, S and some accessory proteins (Figure 2B). Only 13 out of ∼29 proteins of the viral proteome had mutations of significance (Figure 2C). Table 1 lists 27 distinct residues that feature an entropy greater than 0.1 bits, significant in the month of March. In all cases, the incidence of the reference amino acid (ranging 63.6-98.7%) did not increase/decrease more than 18.6%.. However, two contrasting pathways of mutational change were evident in these sites, one in which entropy usually increased and then decreased and the other in which entropy increased constantly (Figure 3):

**Figure 2.**
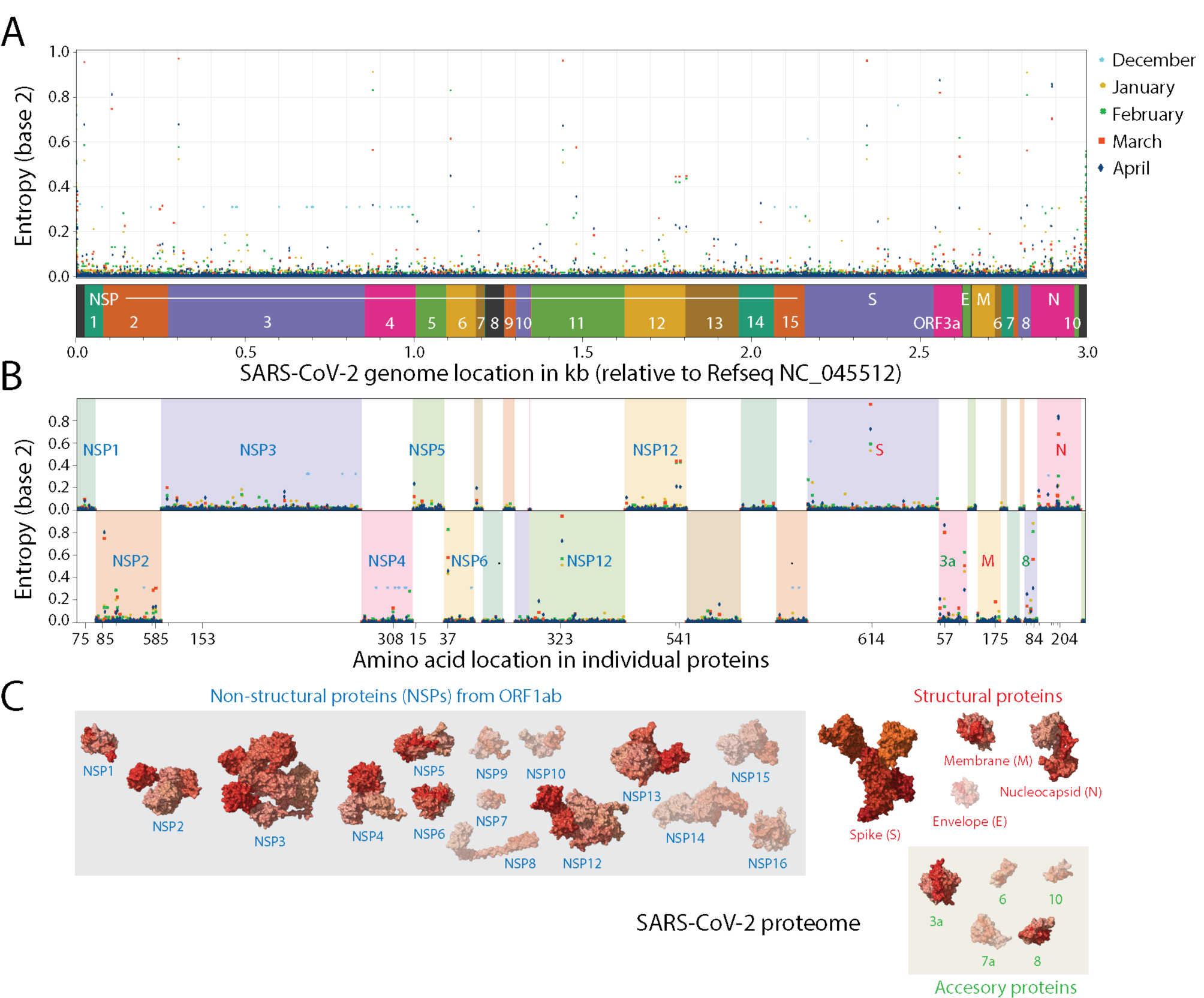
Analysis of mutational entropy at nucleotide (**A**) and amino acid (**B**) levels defines an evolving SARS-CoV-2 proteome of 13 proteins with significant mutational change (**C**). Amino acid locations are only labeled for sites with mutational entropic levels above 0.1 bits in the month of March. Molecules that exhibit significant entropic levels have their atomic 3-dimensional models unshaded in panel C.

**Figure 3.**
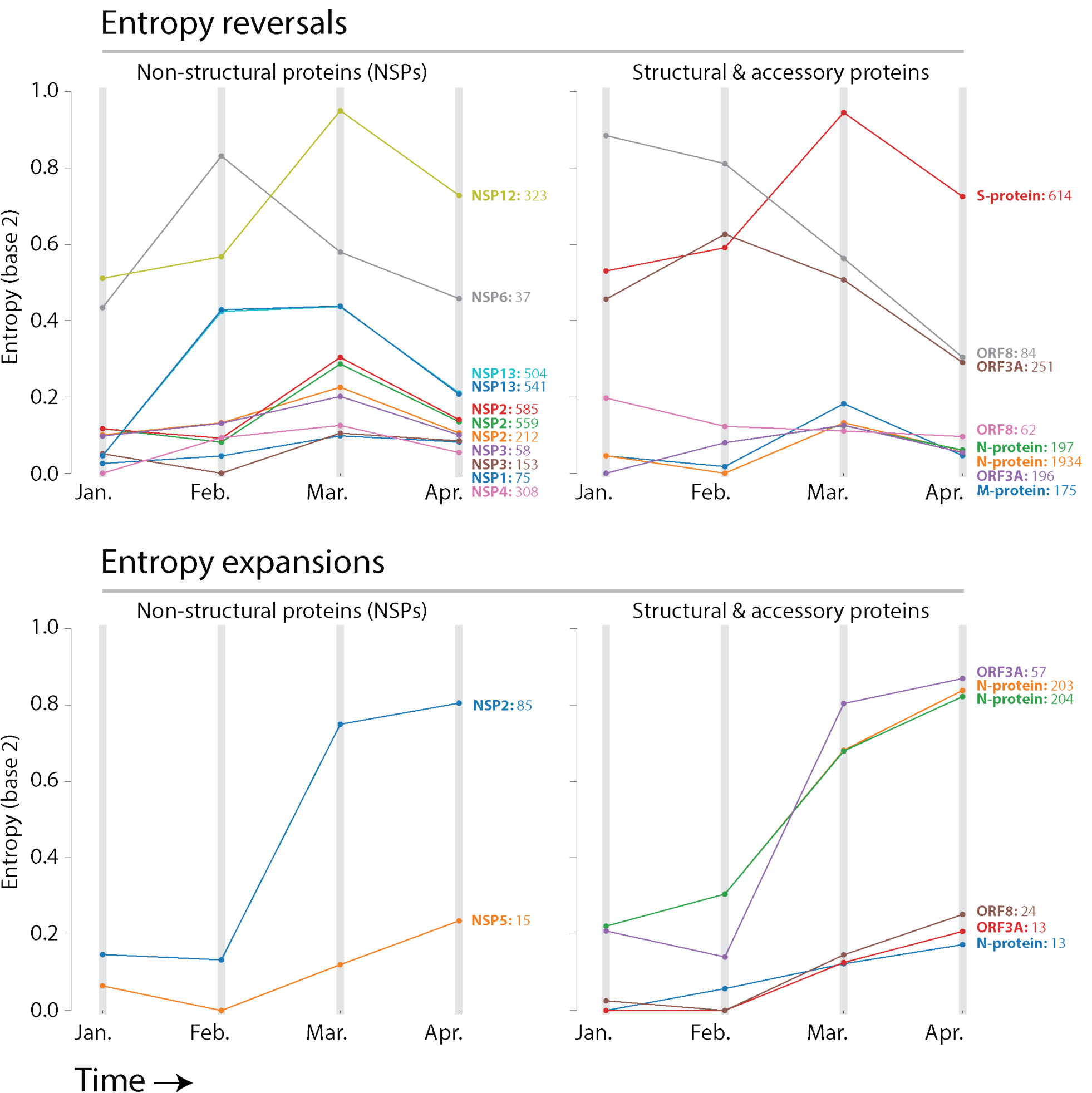
Pathways of mutational change involve mutational entropy reversals and expansions. Entropy reversals occur when entropy increases and then decreases in the timeline of pandemic. Entropy expansions occur when there is only a pattern of increase, which signals continued diversification of amino acid sequences.

i. Entropy reversals: Of the 27 residues with significant entropy, 19 decreased their entropy in April (entropic delta ranging from –0.01 to –0.26). This pattern of entropic reversal is of interest, as a gradually rising entropy followed by a drop suggests the most-recent dominant residue is out-competing the others. Remarkably, the entropy of residues at location 614 of the S-protein of the spike increased continuously from December to March before decreasing −0.22 bits between March and April. However, the continuously increasing incidence of the glycine (G) variant over aspartic acid (D) did not change, rising 15.7% (from 63.6% to 79.33%) between the months of March and April. This observation agrees with previously reported evidence (assessed April 6) of a D614G mutation of S-protein that increases case fatality rate^22^. Similarly, a P323L mutation in NSP12, the viral RNA-dependent RNA polymerase encoded in ORF1b also showed a reversal in entropy of −0.22 bits, as the leucine (L) variant increased in prevalence 18.6% (from 63.8% to 82.4%) over the proline (P) reference, which also decreased 18.6% in incidence. These two locations appear the only sites experiencing significant amino acid swaps. Other sites exhibited less significant patterns. The incidence of leucine (L) of site 84 of ORF8 increased 7.77% (from 86.78% to 94.6%) by directly replacing serine (S). Similarly, the incidence of proline (P) of site 1427 of ORF1b increased 5.63% (from 91.1% to 96.71%) by replacing leucine (L) and the incidence of tyrosine (Y) of site 1476 of ORF1b increased 5.82% (from 90.9% to 96.76%) by replacing cysteine (C). Both mutation sites affect the NSP13 helicase protein (sites p.504 and p.541, respectively). It is important to note that unlike D614G and L323P, the amino acid present in the reference sequence remains dominant in these cases.
ii. Entropy expansions: Eight sites showed entropy constantly increasing but with a concurrent pattern of mutational change (a tendency to swap amino acids) (Table 1, Figure 3). These sites reveal a significant increase in entropy from March to April (entropic delta ranging from 0.05 to 0.16), with a variant residue rising in April with respect to the dominant residue of March. These concurrent tendencies are evident in residues 203 and 204 of the nucleocapsid N-protein. The arginine (R) at site 203 decreased in prevalence 8.33% (from 81.9% to 73.5%) as the incidence of a lysine (K) variant increased 8.2% (from 18.12% to 26.32%). In turn, the glycine (G) residue of the directly adjacent 204 site was exchanged by an arginine (R), increasing 8.2% the incidence of the variant (from 18.1% to 26.3%). In another example, the threonine (T) of site 85 of the NSP2 protein encoded by ORF1a decreased ∼3% in its incidence (from 78% to 75%) as it is replaced by an isoleucine (I) variant. These concurrent patterns of amino acid replacements perhaps indicate that sites will be the next to exhibit a trend of entropy decrease upon reaching a peak.

### Principal Component Analysis

To explore how pathways of mutational change are affecting the viral quasispecies along the timeline of the early pandemic, we performed conventional PCA analysis of genomic samples using BioPython^28,29^ Cluster package, identifying mutation examples of main entropic reversals and expansions (Figure 4). Two distinct elongated clouds described the proteomic make up of evolving viruses, one depicting the departure from the sequence makeup of the reference Wuhan strain (star symbol located on the leftmost part of the cloud) that signals the start of the pandemic and another matching variants in site 614 of the S-protein or sites 203 and 204 of the N-protein. Clouds contained a patchwork of genomes collected throughout the different months of the pandemic, which diversified in sequence along the first component. This is consistent with entropic expansion caused by diversification and drift. However, the temporal incidence throughout the elongated clouds seem to contract towards the location of the reference strain, likely signaling mutational pathways of active fixation are at play. As expected, the genomes harboring the S-protein variant were more numerous than those that preserved the original mutation, which is consistent with the rapid expansion of the D614G mutation and its associated haplotype. In turn, genomes harboring the original N-protein mutations were more abundant, given the slow but steady spread of the variants in the new mutational pathway. We note that the removal of duplicate sequences decreases the variance caused by sequence frequency, causing multidimensional methods to over-represent rare and under-represent common sequences. These are only undesirable if frequency is a variable of interest. As frequency and proportion were already accounted for in the entropy analysis, PCA describes the differences in amino composition, not the likelihood of any given sequence. The inclusion of duplicate sequences would act as noise, skewing the results for the projected dimensions. Regardless of these caveats and justifications, PCA should be merely regarded as a descriptive tool, which satisfactorily performed the role for analysis of the contribution of sequence variances.

**Figure 4.**
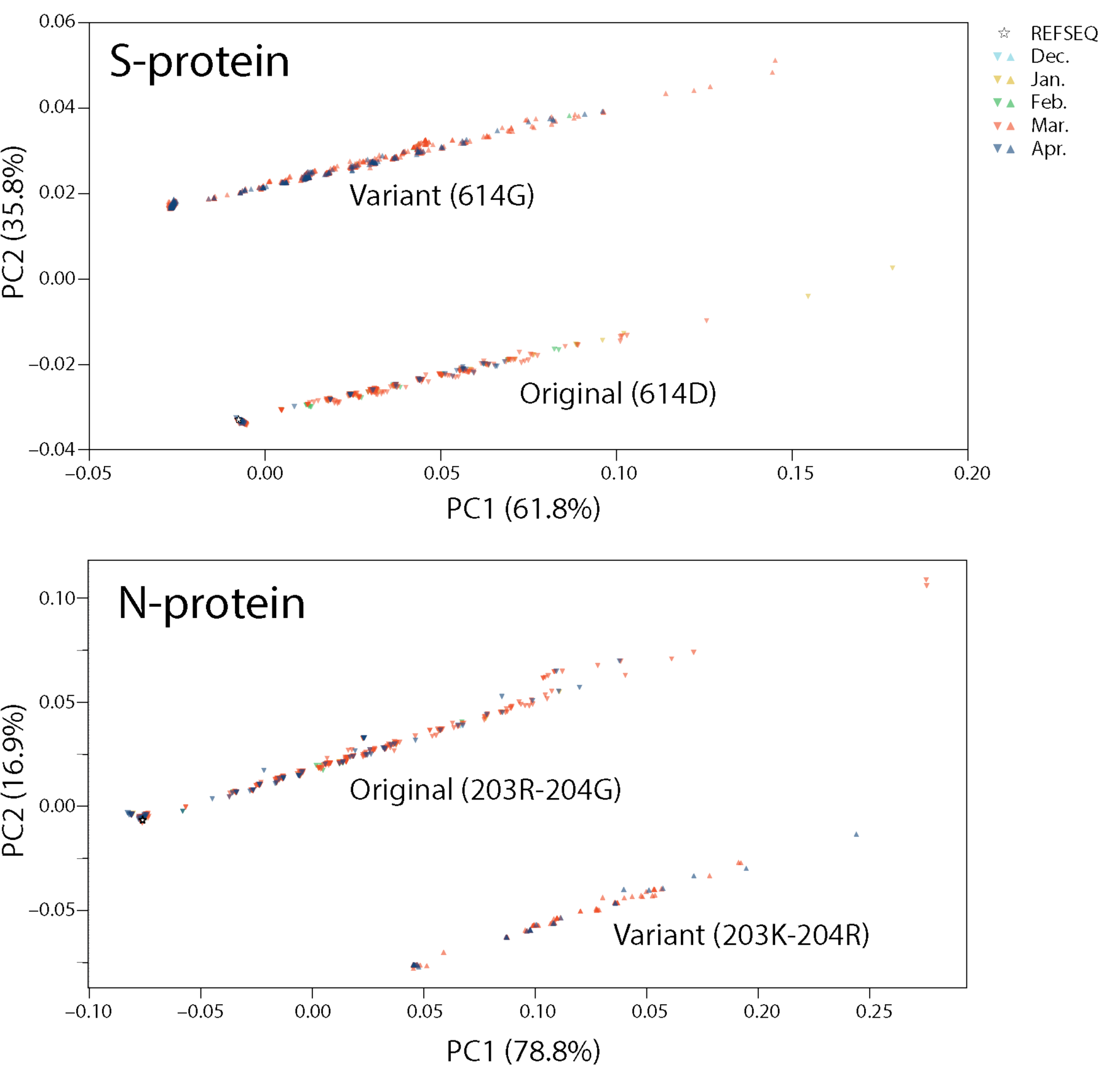
Principal component analysis of SARS-CoV-2 genomes with original and variant sites in the S- and N-proteins.

### Structural Analysis

To make inferences of possible molecular functions affected by the mutants, we traced mutations onto crystal and cryo-EM structures of SARS-CoV-2 proteins and when not available, viral proteins modeled with I-Tasser^32^. Figure S1 describes results. We found that most of the 27 mutation sites were located in loop regions (81%) that were generally on the surface of the molecules (89%). Out of 27 sites, 22 were in loop (also known as turn) regions and 5 in helical regions. No sites were in strand structures. A total of 24 sites were located on the surface of the molecules suggesting an important role in intermolecular interactions. Only two sites were buried (the 203 and 204 mutants of the N-protein) and one faced the pore of accessory viroporin 3a protein. A total of 18 were in ordered regions of the molecules while 7 were in disordered regions (in the N, NSP3 and 3a proteins). Intrinsic disorder and binding propensity scores confirm the disorder of these regions. Sites in NSP4 and protein 3a were located in trans-membrane (TM) regions, consistent with the close association with membranes of these viral proteins.

### Intrinsic Disorder

We performed a global analysis of intrinsic disorder with IUPred2A of the SARS-CoV-1 proteome. It revealed that while intrinsic disorder was variable, the entire protein repertoire was ‘highly structured’. The only notable exceptions were the ‘highly unstructured’ N-protein, the disordered hypervariable (HVR) domain in NSP3, the largest encoded protein of the virus, and disordered terminal regions of viroporins. An analysis of protein intrinsic disorder showed areas exhibiting high disorder scores within the N-protein, indicating significant levels of intrinsic disorder is present in the nucleocapsid (Figure 7). Remarkably, sites 203 and 204 of the N protein that showed increasing entropy and their entropically variable neighborhoods aligned with areas of high disorder. In contrast, sites that experienced no entropic variation (high-conservation) tended to have low disorder. These sites also had high binding site score following an analysis with Anchor2. The conservation of binding sites seems intuitive due to the important functional nature of those regions. In fact, binding site scores peaked around sites 20, 220 and 400 in intrinsic disordered regions and in sites 50, 170, 275 and 355, flanking the two RNA-binding domains (Figure 6). Variable areas of intrinsic disorder raise several possibilities, ranging from being truly disordered functionally unimportant regions to an area experiencing significant functional change.

**Figure 5.**
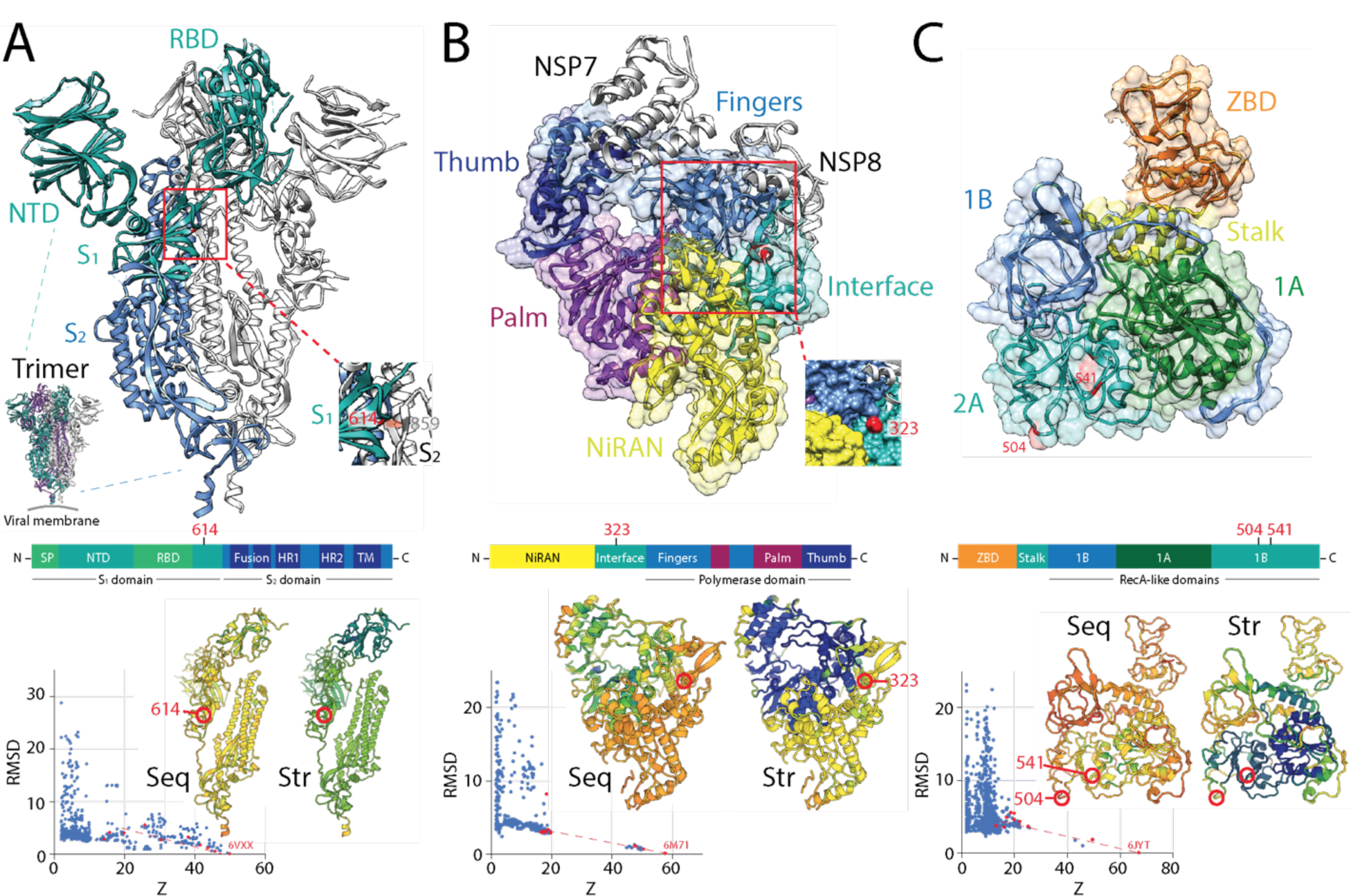
Major SARS-CoV-2 protein molecules experiencing mutational entropic reversals. **A**. The coronavirus spike is a trimer of S-protein protomers, each harboring an N-terminal S_1_ subunit sequence with an N-terminal domain (NTD) and a receptor-binding domain (RBD) and a C-terminal S_2_ subunit holding a ‘fusion’ region with fusion peptide (FD) and internal fusion peptide sequences, two heptad repeat (HR) sequences, and a transmembrane (TM) domain. The subunits are processed by host proteases upon viral entry. The SARS-CoV-2 atomic model of a dimer (PDB entry 6VXX) shows the D614G mutation of the S_1_ domain eliminates a hydrogen bonding interaction with site 859 of the S_2_ domain of another protomer (colored in orange in the inset). An RMSD versus Z-score plot describes the DALI structural neighborhood of 6VXX, which contains 2,310 structures. Alignments of multiple random samples of 10 structures along a transect from 6VXX to the main cloud with low Z-scores of structural similarity (colored red in the plots) consistently show 614 is part of a loop that falls in molecular regions that are poorly conserved at both sequence (Seq) and structural (Str) levels. Blue hues indicate larger conservation than green-to-red hues in the protomer cartoon models of the alignment example. **B**. The NSP12 is the main RNA dependent RNA polymerase of the virus. It is encoded by ORF1b and is responsible for the synthesis of viral RNA. Examining the SARS-CoV-2 structure in complex with NSP7 and 8 cofactors (PDB entry 6M71) revealed that the P323L mutation sits in an ‘interface’ region (spanning residues 250-365) between the N-terminal nidovirus-unique NiRAN domain with nucleotidyltransferase activity and the C-terminal polymerase domain that harbors the fingers, palm and thumb subdomains. The mutation is in a helical region at the surface of a pocket formed by the NiRAN, interface and fingers structures (inset). DALI structural neighborhood analysis (1,157 structures) confirmed the site is in a region that is poorly conserved at sequence and structure levels, but borderline with the highly conserved regions that harbor polymerase activity.**C**. The NSP13 is the helicase of the viral replication complex. NSP13 has an N-terminal Zn binding domain (ZBD) followed by a stalk domain and three Rec-A domain structures 1A, 2A and 2B, which form the triangular base of a pyramid. L504P and C541Y are in loop regions located on the surface of the middle of the 2B domain. DALI structural neighborhood analysis of PDB entry 6JYT (1,356 structures) showed C541Y is in regions of the molecule that are structurally conserved, while the L504P region was variable at both sequence and structure levels.

**Figure 6.**
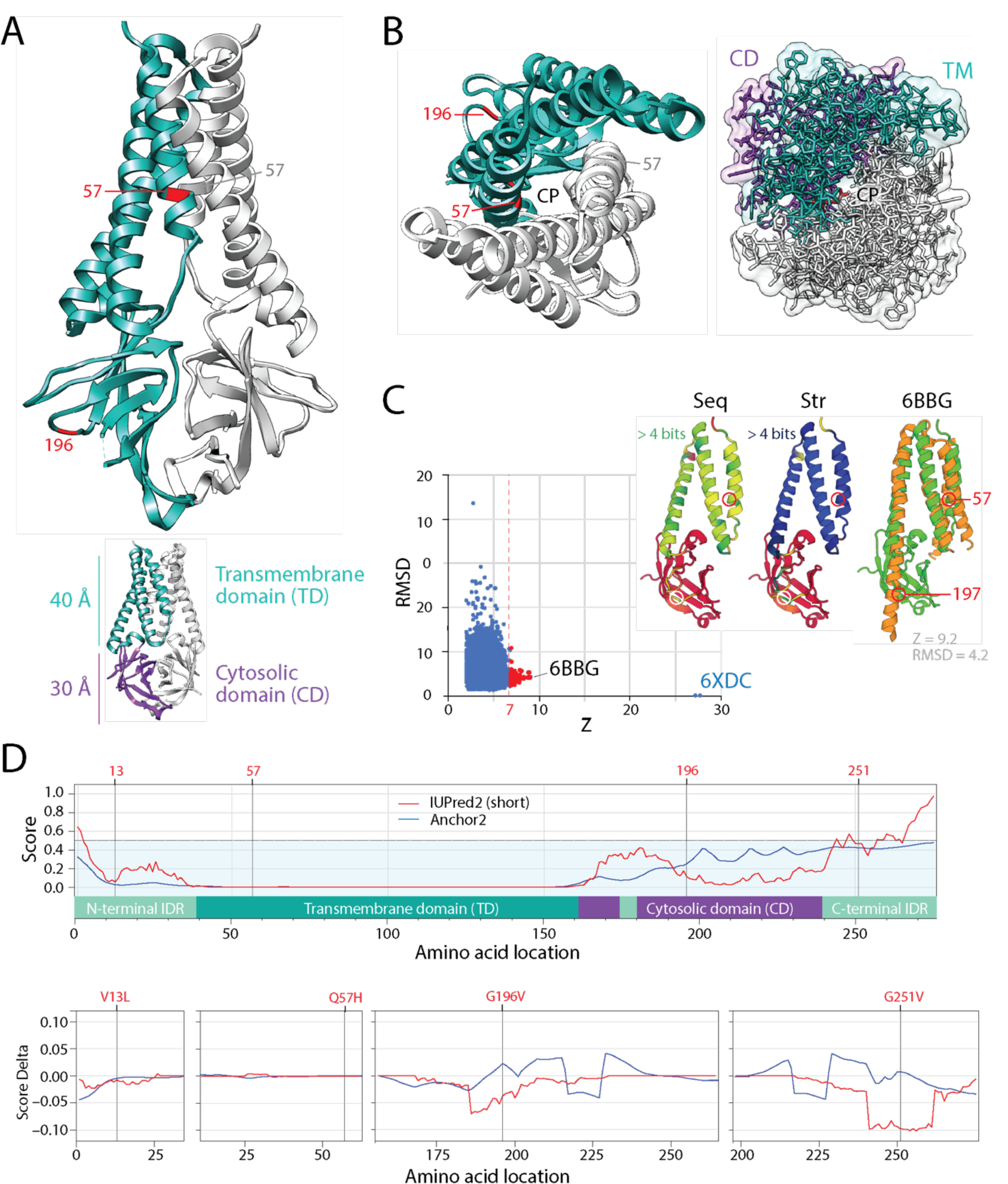
The mutational diversification of SARS-CoV-2 viroporin encoded by ORF3a. **A**. The structure of the protein 3a molecule (PDB entry 6XDC) has two domains, a N-terminal transmembrane domain (TD) and a C-terminal cytosolic domain (CM). Mutation Q57H is located in the first of the three transmembrane helices at the major hydrophilic constriction of the pore important for channel activity. Mutation G197V forms part of a loop at the surface of the CD. Terminal amino acids 1-38 and 239-275 and a 175-180 in CD could not be modeled because they were weakly resolved. They hold mutations 13 and 251. **B**. View from the lumen side of the channel pore (P) in ribbon and atom stick representation. Note that the pore is only 1 Å wide. **C**. A DALI structural neighborhood analysis (10,088 structural neighbors) returned significant hits to small fragments (Z ≤ 9.2; RMSD ≥ 1.3) that formed a single cluster in the RMSD versus Z-score plot. Structural alignment of the 92 hits with Z ≥ 7 (red dots) revealed that all hits matched the TD structures and were well conserved at structure (Str) but less at sequence (Seq) levels. The best structural match to the TD was the Orai protein channel (PDB entry 6BBG) responsible for Ca^2+^ influx pathways in metazoan cells and involved in immune responses and cancer. **D**. The mapping of intrinsic disorder (UIPred2, red line) and gain-loss of binding energy (Anchor2, blue line) along the sequence confirmed the significant intrinsic disorder (scores ≥ 0.5) of the C-terminal linker. A comparison of the different mutants and reference viral strain with a delta score revealed that mutations G196V and G251V decreased disorder.

**Figure 7.**
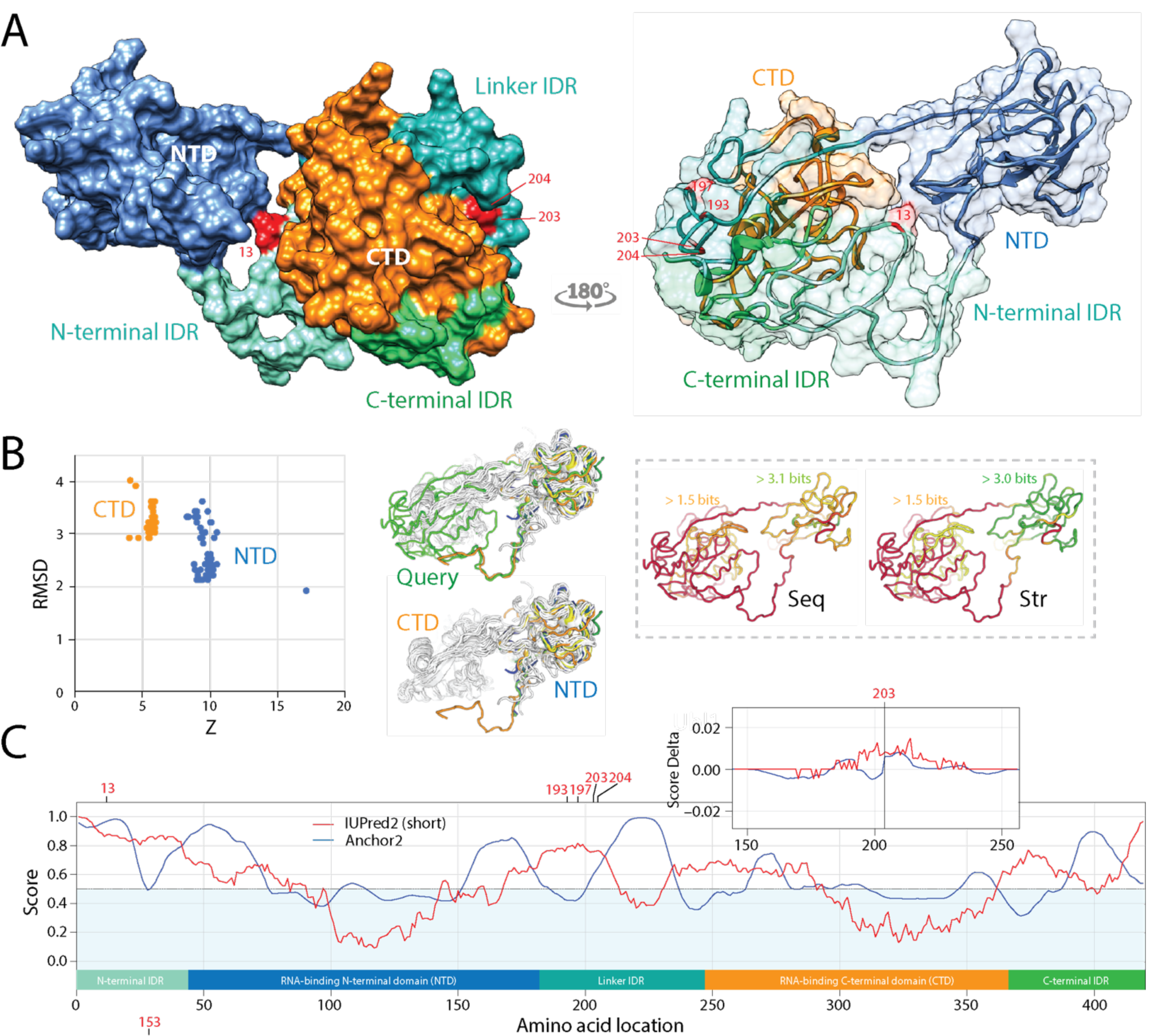
Pathways of mutational diversification of SARS-CoV-2 involve intrinsic disordered regions of the nucleocapsid (N) protein. **A**. The N-protein has two major RNA-binding domains, an N-terminal domain (NTD) and a C-terminal domain (CTD), both connected to a central linker and flanked by terminal sequences, all of which have been reported to be intrinsically disordered regions (IDRs). Mutations were traced onto a SARS-CoV-2 N-protein structure modeled with I-Tasser. They occurred in position 13 of the N-terminal IDR and positions 193, 197, 203 and 204 of the linker IDR, all of them in loop regions of the molecule. Mutations 203 and 204 were the only sites that were buried in the molecule. **B**. A DALI structural neighborhood analysis against the modeled structure (88 structural neighbors, including many from SARS-CoV-2) showed two clusters in the RMSD versus Z-score plot, one reflecting structural match to the NTD domain and the other to the CTD domain. Structural alignment plots of the 88 structures supported the veracity of the modeled RNA-binding domains and revealed that the NTD is more conserved at sequence (Seq) and structure (Str) levels. **C**. The mapping of intrinsic disorder (UIPred2, red line) and gain-loss of binding energy (Anchor2, blue line) along the sequence confirmed the significant intrinsic disorder and binding (scores ≥ 0.5) of linker and terminal regions. A comparison of the R203K mutant and reference viral strain with a delta score revealed that the mutation increased disorder. A similar outcome was obtained with the G204R mutant.

## Discussion

RNA viruses are known for their high mutation rates, and hence, for rapid genome evolution^39^. One consequence of these rates is that viral populations become ‘quasispecies’, highly diverse collectives of closely related viruses expressing vast numbers of distinct genotypes^40^. Coronaviruses harbor the largest known non-segmented RNA genomes reported to date (∼27-32 kb in length) and a proteome with a repertoire of ∼29 proteins, many with published atomic structures (Figure 2). They adapt to host environments by relying on the low fidelity of a replication complex, which assembles around the NSP12 polymerase with its extra N-terminal β-hairpin domain in interaction with an hexadecameric complex of disordered NSP7 and NSP8 cofactors that are currently targets of COVID-19 therapeutics (e.g. *remdesivir* postinfection treatment)^41,42^. While the limited proofreading and repair capabilities typical of RNA viruses puts coronaviruses close to an ‘error threshold’ of too many deleterious mutations for successful persistence^40^, their proofreading is crucially enhanced by the 3’-to-5’ exonuclease domain of the NSP14 protein^43^. This additional enzymatic activity and interaction increases 12-15-fold the template copying fidelity of coronaviruses^44^, which protects them from the error threshold relationship and enables both genome expansion and mutation robustness^39^. It also allows for an ample mutant spectrum fueled by the average mutation rates of ∼1 mutation per genome per round or replication that are typical of RNA viruses^45^. Indeed, our study of 12,606 evolving genomic sequences finds the expected complex mutant spectrum responsible for pathways of mutational change operating in SARS-CoV-2 populations during the initial stages of the COVID-19 pandemic. These pathways cannot be considered neutral traits. As we will now discuss, pathways specifically involve the replication complex and innate and adaptive immune responses triggered by the spike, the nucleocapsid, and other proteins, all of which confer selective advantage to the viral quasispecies. Interestingly, while we find highly entropic sites in the NSP14 exonuclease protein at nucleotide level (Figure 2A), we identified no significant entropic diversity at the amino acid level in this crucial proofreading molecule (Figure 2B, Table 1). This suggests strong selective pressure to maintain invariant the NSP14 amino acid sequence, possibly because its exonuclease activity is central for viral quasispecies diversity.

Most entropically-significant mutations in NSPs showed entropy ‘reversals’, in which entropy increases were followed by decreases along the timeline of the pandemic (Figure 3). In turn, mutations in structural and accessory proteins followed both modes, entropy reversals and entropy expansions. We first illustrate entropy reversals with three high-entropy mutations, one in the structural protein of the coronavirus spike and the other two with mutations in NSP proteins important for virus replication:

### S-protein

As previously reported^22,28^, our research exploration reveals the active fixation of the D614G mutation of the S-protein of the spike, which we also find has coordinated entropic trends associated with the P323L mutation of the NSP12 polymerase that mediates viral replication (Figure 3). Note that D614G is part of a haplotype of 4 mutations (including those that alter NSP12, the 5’ UTR, and silently NSP3), which constitute the G-clade that originated in China and was established in Europe^28^. *In silico* modeling suggests the G614 variant of the spike: (i) breaks a D614 −T859 side chain hydrogen bond between the neighboring S_1_ and S_2_ units of different spike protomers, thereby enhancing flexibility and modifying their interactions,^22,28^ (ii) modulates glycosylation of the neighboring N616 site^28^, and/or (iii) alters the dynamics of conformational transitions of the proximal fusion peptide^28^. Figure 5A traces the location of the mutation in an atomic model of the spike of SARS-CoV-2 and reveals it falls in regions of the molecule that are poorly conserved at both sequence and structure levels. Remarkably, the G614 variant was found associated with increased viral loads and higher infectivity, but not disease severity^28^. However, in another study the variant was shown to be strongly correlated with case fatality rate^22^. While the link to virulence needs confirmation, the coronavirus S-protein mediates entry into host cells and the residues within the RBD are critical for determining virus transmission and host range^16^. In an earlier study, several mutations in the RBD were found to enhance SARS-CoV (K479N and S487T) and SARS-CoV-2 (493 and 501) interaction with human ACE2 binding hot spots Lys31 and Lys353, and mutations K479N and S487T played a critical role in the civet-to-human and human-to-human transmission, respectively^16,19^. Ferrets, cats, pigs, orangutans, monkeys share similar critical virus-binding residues with humans, suggesting SARS-CoV-2 could infect these animals^16^. None of these sites were significantly mutated in the expanding SARS-CoV-2 populations we sampled, stressing the fact that the current pandemic is mediated by human transmission through stabilized binding capacity to human ACE2. However, future mutations could expand transmission potential and host range to for example domesticated animals (e.g. cats, ferrets, and hamsters, which are infected and spread the disease)^46^.

### NSP12 polymerase

As part of the 4-mutation haplotype, the P323L mutation of the NSP12 polymerase is also actively fixed (Table 1). NSP12 contains a nidovirus-unique N-terminal extension domain with a nucleotidyltransferase (NiRAN) architecture that is connected to the C-terminal polymerase domain (with its fingers, palm and thumb subdomains) through an ‘interface’ domain that spans residues 250-365^41^. P323L seats at the center of the interface domain, contacting the NiRAN and fingers domains of NSP12 and the second subunit of NSP8 (Figure 5B). The mutation site is at the surface of a pocket formed by the NiRAN, fingers, and interface regions that is poorly conserved at the sequence and structure levels. The interface domain acts as a protein-interaction junction^42^.

### NSP13 helicase

One novelty is that our analysis shows active fixation tendencies of two mutations in the NSP13 helicase protein that mediates viral RNA unwinding, L504P and C541Y (Figure 5C). The NSP13 helicase catalyzes the NTP-dependent unwinding of oligonucleotide duplexes into single strands. NSP13 adopts a triangular pyramid shape structure with 5 domains in SARS-CoV^47^ and in modeled SARS-CoV-2 molecules^48^. Three Rec-A domain structures, 1A and 2A and 2B form a triangular base for the N-terminal Zn binding domain (ZBD) and the stalk domain. The triangular base forms a nucleic acid binding channel with 1A crucially participating in unwinding, in which 1A tightens the grasp on the nucleic acid while 2A loosens the grip in each translocation unwinding step^47^. The sites of the L504P and C541Y mutations fall on the N-terminal region of the 2A domain and are located in loops at the surface of the domain structure (Figure 3C). These mutations are therefore center stage for translocation functions. In addition, helicase translocation structure 1A and the ZF3 motif of the ZBD domain interact with the NSP12 polymerase during viral replication^47^. The close polymerase-helicase interaction could explain the joint fixation of the NSP12 and NSP13 mutations we have identified.

The coordinated reversals of entropic expansions provide support to the fixation of mutations in the spike, polymerase and helicase proteins, some of which fostered virulence and viral loads. A number of mutations that we do not discuss followed this same entropic mode but achieved lower entropy levels (Table 1; Figure 3). For example, the large NSP3 scaffold and protease that initiates cleavage of the ORF1a/ORF1ab protein and mediates genome replication and transcription^11^, harbors a mutation at the hypervariable region (HVR), which decreases disorder in only the disordered domain of the protein (Figure S2). While HVR appears dispensable for viral infection, its role is currently unknown^11^. In parallel, we also identified significant entropic regions with persistent tendencies of entropic expansion that could be important determinants of disease progression (Table 1; Figure 3). Here we focus on high-entropy mutations affecting two important molecules, the accessory viroporin 3a protein and the N-protein, which we posit represent new pathways of mutational change that involve intrinsic disorder:

i. Viroporin protein 3a: Viroporins are small hydrophobic ion-channel proteins that modify cellular membranes to facilitate virus release, replication and virulence^49^. Coronaviruses have three viroporins, the E-protein and accessory proteins 3a and 8, the first two required for viability and virulence^50^. Protein 3a is the largest of the three viroporins (Figure 2C) and is encoded by ORF3a. The cryo-EM structure of the SARS-CoV-2 viroprotein has been recently acquired in lipid nanodiscs^51^. Protein 3a molecule has two domains, a N-terminal transmembrane domain (TD) and a C-terminal cytosolic domain (CM) with a new globular fold (Figure 6A). The 40 Å high TD region has 3 helices per protomer, with their N-termini oriented towards the lumen. Viewed from the lumenal (extracellular) side (Figure 6B) the 6 helices of two promoters arrange in clockwise order along an elliptical trajectory. The first two and the last two helices are joined by short intracellular and extracellular linkers, respectively. The last helix of the TM domain connects to the globular cytosolic CD via a helix-turn-helix motif. A DALI structural neighborhood analysis of the cryo-EM structure returned 92 significant hits to small fragments of Z ≥ 7 (red dots in the RMSD vs Z plot), all of which aligned to the TM region (Figure 6C).Their alignment revealed matches to the structures of the TD region that were well conserved at structure but less so at sequence levels. The best structural match was to transmembrane units of the Orai protein channel responsible for store-operated Ca^2+^ influx pathways in metazoan cells^52^. The Orai proteins are necessary to activate immune responses and are involved in a vast range of physiological processes important for cancer research. It is therefore likely that protein 3a facilitates virus spread by helping dissipate membrane potential for cell lysis and virus release using store-operated mechanisms similar to those of Orai proteins. The high-entropy mutation Q57H, which follows an entropy expansion mode (Figure 3), is located in the first transmembrane helix at the major hydrophilic constriction of the pore, which is important for ion channel activity. However, the mutation did not affect the main function of the ion channel, perhaps because the mutated residue does not affect ion channel properties^51^. In turn, low-entropy mutation G197V, which follows the entropic return mode (Figure 3), forms part of a loop at the surface of the CD region. Mutations 13 and 251 are located in terminal amino acids 1-38 and 239-275 regions, respectively. Low entropy mutation 13 follows an entropy expansion mode while high-entropy mutation 251 follows an extropy return mode. Together with a 175-180 amino acid region in CD, these regions could not be modeled because their cryo-EM structures were weakly resolved. Kern et al.^51^ suggested these regions of protein 3a represented areas of intrinsic disorder. Proteins are intrinsically flexible molecules. While structural flexibility is a property of regular (e.g. helices, strands) or irregular (e.g. loops) structure that is necessary for the establishment of molecular functions, proteins can also exhibit disorder, lack of significant constraints on internal degrees of freedom of the polypeptide chain. Intrinsically disordered regions are flexible regions of proteins that lack a fixed 3-dimensional (3D) structure under native cellular conditions but are generally involved in a variety of molecular functions^53^. These regions usually apply to protein backbones that lack consistent Ramachandran dihedral angles, i.e. they are dynamic and exhibit conformations that resemble either random-coils, molten globules or flexible linkers. We mapped intrinsic disorder and gain-loss of binding energy along the sequence of the protein 3a viroprotein (Figure 6D). Intrinsic disorder levels classified the protein as ‘moderately disordered’ according to the fraction of residues that were unstructured^34^. However, the analysis revealed highly significant intrinsic disorder (scores ≥ 0.5) were present in the C-terminal and to a lesser degree in the N-terminal regions of the molecule, supporting the hypothesis that these regions are not resolved in the cryo-EM density map because of disorder. In addition, a comparison of mutants V13L, Q57H, G196V and G251V and the reference viral strain with delta scores of disorder and binding revealed interesting patterns. Mutations in the N-terminus and the pore did not alter significantly both disorder or binding. In contrast, G196V in the loop region of CD and G251V in the C-terminus region induced significant decreases in disorder and increases in binding potential in regions as far away as 30-50 amino acids from the mutated residue. The long-range effect and magnitude of these mutations is significant, especially G251V, which is present in 13.8% of all proteomes examined and its entropic incidence is growing.
ii. N-protein: We also identified pathways of mutational change in the N-protein that enhance protein intrinsic disorder and affect viral spread. The N-protein plays critical roles in maintaining viral structure and viability once the virus has entered the cell, including replication and transcription, and packaging of the virus RNA^23^. The N-protein has two major RNA-binding domains, an N-terminal domain (NTD) and a C-terminal domain (CTD), both connected to a central linker and flanked by terminal sequences, all of which have been reported to be intrinsically disordered regions (IDRs). Crystal structures of the SARS-CoV-2 NTD [PDB entries 6M3M^54^ and 6WKP^55^] and CTD [6WJI^56^] have been recently deposited. The two domains align well to a complete SARS-CoV-2 N-protein structure modeled with I-Tasser (data not shown), enabling its use to trace mutations in the IDRs of the model (Figure 7A and B). A DALI structural neighborhood analysis against the modeled structure (88 structural neighbors, including many from SARS-CoV-2) showed two clusters in the RMSD versus Z-score plot, one reflecting structural match to the NTD domain and the other to the CTD domain. Structural alignment plots supported the veracity of the two modeled RNA-binding domains and revealed that the NTD is more conserved at sequence and structure levels. The two domains of the N-protein are heavily phosphorylated, which favors binding to the viral RNA genome in a beads-on-a-string type conformation but with different mechanisms^57^. Indeed, we find that Anchor2 binding propensity is significant at the N-terminal regions of the NTD and CTD regions (Figure 7C). The N protein also binds the NSP3 component of the replicase-transcriptase complex and the M-protein. These multiple interactions likely help package the encapsidated genome into viral particles. Intrinsic disorder levels classified the N-protein protein as ‘highly disordered’ and this property manifested in the panel of mutations identified. Mutations occurred in position 13 of the N-terminal IDR and mutations in positions 193, 197, 203 and 204 occurred on the central linker IDR, all of them in loop regions of the molecule (Figure 7A). Mutations 203 and 204 were the only sites that were buried in the molecule. We identified significant increases in the entropy of residues 203 and 204 of the N-protein, which reached Shannon entropy values of ∼0.8 (Figure 3). This finding suggests an active and ongoing quasispecies exploration for novel genotypes. The coordinated rates of increase of the R203K and G204R mutations indicated mutations did not occur randomly. Instead, functional constraints appear to act on the mutation process suggesting entropic trends will continue until dominant genotypes become fixed in the viral population. The fact that all significant mutations occurred in IDRs of the N-protein suggests intrinsic disorder plays a biological role in the spread of these mutations. This is not surprising since intrinsic disorder mediates coronavirus transmission. The unstructured disorder of M and E proteins appear responsible for the rigidity of the coronavirus shell and the transmission mode (respiratory or orofecal tropism) of the virus^58^. SARS-CoV-2 retains the flexibility of the respiratory mode while enhancing the environmental resilience of the virion for efficient dispersion^59^. To confirm the role of intrinsic disorder, we mapped disorder (UIPred2, red line) and gain-loss of binding energy (Anchor2, blue line) along the sequence (Figure 7A). Indeed, significant disorder and binding levels (scores ≥ 0.5) existed in linker and terminal regions. The R203K and G204R mutations spanned regions of high disorder and high binding ability. Both mutations increased intrinsic disorder relative to the reference Wuhan strain (Figure 7C). A functional role of this highly disordered binding region is expected, including the interaction with a wide variety of targets that is considered a hallmark of intrinsic disorder^60^. While the role of the amino acid changes within the N-protein region remains biologically uncharacterized in our investigations, mutations suggest the virus is attempting to optimize replication inside the host cells by tuning both flexibility and binding.

Our study explores pathways of mutational change in highly entropic sites of the SARS-CoV-2 proteome, quantifying diversity and fixation of variants with the two-state variable strategy we modified from Pan and Deem^38^. The analysis does not explore phylogenetic relationships at genomic level with distance or parsimony-based reconstruction methods, which for example are systematically pursued in the Nextstrain portal^27^. Instead, we recognize the difficulties of studying the multidimensional landscape of rapidly evolving and recombinogenic coronavirus genomes with tree-based approaches that do not dissect processes of horizontal exchange of genetic information, do not model the short-timescale accumulation of mutations, and are poorly powered to resolve the existence of positive selection. Instead, we trace mutation accumulation in the expanding SARS-CoV-2 population of the pandemic to uncover significant pathways of proteome diversification. A conventional PCA analysis conducted on a distance matrix comprised of distinct amino acid sequences, excluding duplicates, revealed a complex data structure with two distinct diversification clouds unfolding fundamentally on the first dimension as new proteome sequence variants depart from the makeup of the original Wuhan reference virus. These two clouds reflected the two mayor entropic pathways of change we uncovered in our analysis.

## Conclusions

A number of important mutations that foster viral spread, such as the haplotype that affects the S-protein, follow an entropic reversal mode that suggest they are being actively fixed in the expanding viral population of the pandemic. Here, we describe new pathways of entropic expansion of mutations involving intrinsically disordered regions of the SARS-CoV-2 proteome and interactions with replicating genomes and endoplasmic membranes that are needed for virus assembly and release from infected cells. Pathways involve intrinsically disordered regions of the N-protein, a structural protein that forms complexes with genomic RNA, interacts with the M-protein during virus assembly, enhances the efficiency of virus transcription and assembly, and help overcome the host innate immune response^61^. Pathways also involve viroporins. Viroporins of enveloped viruses such as SARS-CoV-2 insert into membranes to break chemoelectrical barriers by channeling ions across membranes and dissipating membrane potential, a property that stimulates budding and resembles that of depolarization-dependent exocytosis^49^. The close homology of SARS-CoV-2 protein 3a to the Orai proteins suggests membrane potential dissipation involves store-operated channeling mechanisms that control Ca^2+^ cellular levels. These new mutational pathways may be responsible for new symptomatic manifestations of the COVID-19 disease. For example, asymptomatic SARS-CoV-2 infection has been reported to account for ∼50% of total coronavirus cases and the majority of the patients with mild infection symptoms can recover by themselves^62,63^. In turn, the COVID-19 disease mechanism suggests that the severe symptoms of COVID-19 involve the uncontrolled immune response of the host^64^. Indeed, mutational pathways involve virus molecules that can subvert the immune response, specifically the interferon response. For example, the N-protein, protein 3a and accessory protein 6 are the three β-interferon antagonists operating in coronavirus disease^65^. Mutations in two of these molecules are high entropy in our mutational set. In addition, the SARS-CoV-2 new virus tends to be more rapidly spreading and less lethal than SARS-CoV and MERS-CoV, a fact that demands explanation. As predicted by the director of Centers for Disease Control and Prevention of the USA on March 25, *“This virus is going to be with us. I am hopeful that we will get through this first wave and have some time to prepare for the second wave*.*”* We hope the exploration of mutational pathways can anticipate moving targets for speedy therapeutics and vaccine development as we prepare for the next wave of the pandemic.

## Acknowledgements

This study began as a class research project in CPSC 567, a course in bioinformatics and systems biology taught by G.C.-A. at the University of Illinois in the spring of 2020. T.T., R.S.D and M.D. conceived the experiments and analyzed the data, with help from G.C.-A. All co-authors wrote and edited the manuscript. We dedicate this work to the frontline medical professionals who have been saving the life of others with limited protective equipment, selflessly, and at their own peril. We also thank public health professionals and scientists for making real-time data and sequences readily accessible to the public.

---

**Figure S1.**
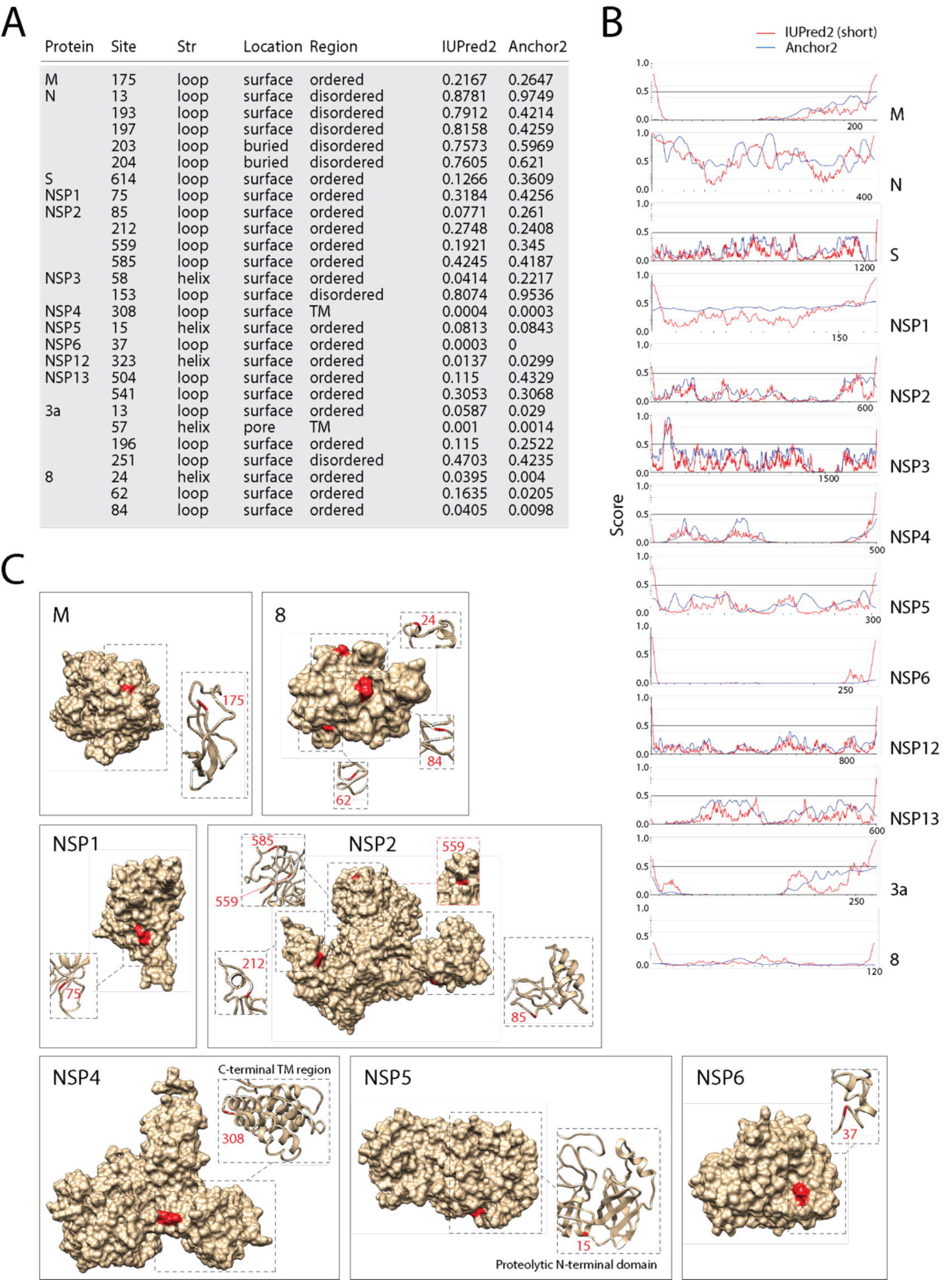
Structural analysis of the SARS-CoV-2 proteome with significant mutational hits. **A**. An examination of the structure (Str: loop, helix, strand), location (surface, buried, pore) and region [ordered, disordered, transmembrane (TM)] of the mutation sites are provided together with scores of intrinsic disorder and binding propensity. **B**. Intrinsic disorder and binding propensity analysis along the sequence of individual proteins with significant entropic sites. **C**. Structural models show structure and location of mutation sites highlighted in red. Missing proteins can be found in Figures 5. 6, 7 and S2.

**Figure S2.**
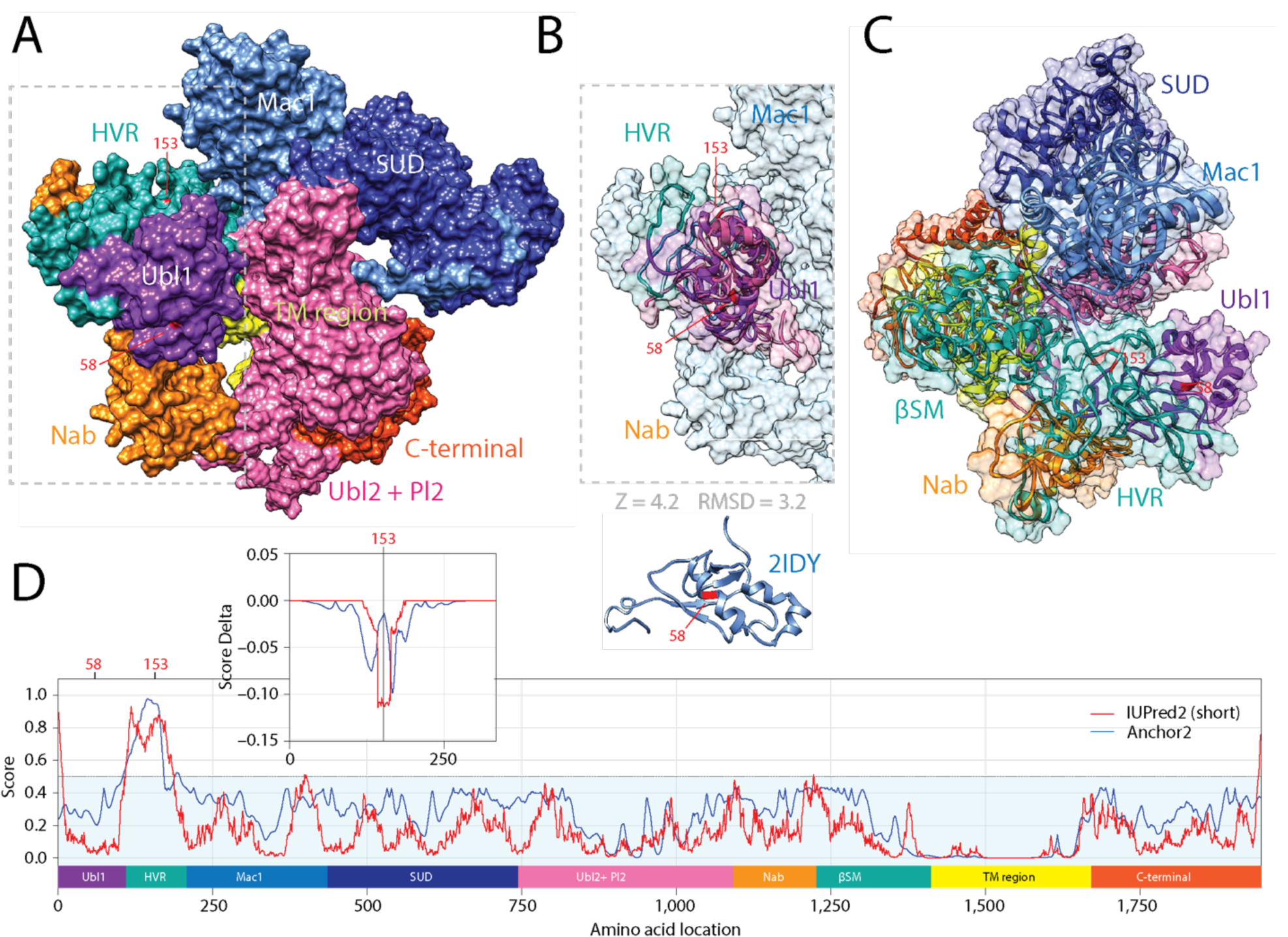
The mutational diversification of SARS-CoV-2 NSP3 involves the only intrinsic disordered domain of the large multidomain scaffold–protease protein. **A**. Mutations at positions 58 and 153 were traced onto the SARS-CoV-2 NSP3 structure modeled with I-Tasser and depicted as surface representation in Chimera. Both were located in crevices of the surface. The large protein contains 16 domains, including the ubiquitin-like domain 1 (Ubl1), the hypervariable region (HVR) acidic domain, macrodomain 1 (Mac1), a SARS-unique domain (SUD) composed or two macrodomains and DPUP, ubiquitin-like domain 2 (Ubl2) and the papain-like protease 2 domain (Pl2) that cleaves the protease from ORF1a/ORF1ab, the nucleic acid-binding domain (Nab), the β-coronavirus-specific marker domain (βSM), a transmembrane (TM) region and a C-terminal multidomain region. **B**. Structural alignment of the Ubl1 domain (PDB entry 2IDY) to the modeled structure reveals statistical significant structural matches. Mutation 58 is highlighted in red on the aligned structures and on the 2IDY model. **C**. A ribbon representation of the modeled structure shows that mutation 58 is located in a helical region of Ubl1 and mutation 153 in a loop region of HVR, which is known to be intrinsically disordered. **D**. The mapping of intrinsic disorder (UIPred2, red line) and gain-loss of binding energy (Anchor2, blue line) along the NSP3 sequence confirmed the significant intrinsic disorder and binding (scores ≥ 0.5) of the HVR region. No intrinsic disorder was observed in the βSM region, despite reports of its intrinsic disorder in other the β-coronavirus sequences. A comparison of the L153P mutant and reference viral strain with a delta score revealed that the mutation decreased disorder.

